# Towards quantitative DNA Metabarcoding: A method to overcome PCR amplification bias

**DOI:** 10.1101/2023.10.03.560640

**Authors:** Sylvain Moinard, Didier Piau, Frédéric Laporte, Delphine Rioux, Pierre Taberlet, Christelle Gonindard-Melodelima, Eric Coissac

## Abstract

Metabarcoding analyses have recently undergone significant development due to the power of this technique in biodiversity monitoring. However, it is still difficult to draw accurate quantitative conclusions about the ecosystems studied, mainly because of biases inherent in the environmental DNA or introduced during the experimental process. These biases alter the relationship between the amount of DNA observed and the biomass or number of individuals of the species detected. Two of the biases inherent in metabarcoding have been measured: the ratio between total DNA and target DNA concentrations, and the PCR amplification bias. A method for their correction is proposed. All experimental tests were performed on mock alpine plant communities using the marker *Sper01*, which is expected to have low amplification bias due to its highly conserved priming sites. Our approach combines standard quantitative PCR techniques (qPCR and digital droplet PCR) with a realistic stochastic model of PCR dynamics that accounts for PCR saturation. The model was used to estimate PCR efficiencies for each species and to infer the true species proportions of the mock communities from the read relative frequencies. The corrections are easy to implement and can be applied to previously generated DNA metabarcoding data. This work demonstrates the relative importance of the two biases considered and is an open door to quantitative metabarcoding data, although many other biases remain to be considered.

## Introduction

In the context of mass species extinction (Barnosky et al., 2011), biodiversity assessment is currently a major challenge. Classically, biodiversity inventories consist not only of a list of species occurring at a site, but also of quantitative data assessing the abundance of each species. Traditional approaches based on direct observation by taxonomists may be unrealistic in terms of available skills and costs, given the enormous effort required to conduct such a survey on a global scale and across the tree of life. Therefore, high-throughput methods, including DNA metabarcoding (Taberlet, Coissac, Pompanon, Brochmann, & Willerslev, 2012), are the only chance to achieve such a goal. DNA metabarcoding has been used for more than a decade in many areas of ecology, such as biodiversity monitoring (*e.g*. Bohmann et al., 2014), detection of invasive species (*e.g*. Klymus, Marshall, & Stepien, 2017), or tracking animal diets (*e.g*. Pompanon et al., 2012). It is now part of the basic toolbox of ecologists, if we consider more than a thousand articles published annually based on this technique. While metabarcoding provides a satisfactory species inventory (Beng & Corlett, 2020; Ficetola & Taberlet, 2023; Taberlet et al., 2012) with some insight into their relative abundance (Pornon et al., 2016), the quality of quantitative data produced is questionable (Krehenwinkel et al., 2017; Yang et al., 2021).

The relationship between the abundance of a species in the field and the number of sequence reads measured in a DNA metabarcoding experiment is far from straightforward. Many reasons can lead to biased abundance estimates. Biases arise from both natural properties and technical issues (Luo, Ji, Warton, & Yu, 2022; van der Loos & Nijland, 2021). At least three natural biases can be considered. First, if the amount of DNA shed into the environment depends on the biomass of individuals (Elbrecht & Leese, 2015; Elbrecht, Peinert, & Leese, 2017; Lamb et al., 2019), it is also a function of shedding rates specific to each DNA source (Wilder, Farrell, & Green, 2023). Second, the relationship between the eDNA sampled, and the DNA actually shed depends on its decay rate, which in turn depends on the ecosystem studied (Andruszkiewicz Allan, Zhang, Lavery, & Govindarajan, 2021; Krehenwinkel et al., 2018). Third, the number of copies of the DNA marker targeted by metabarcoding per unit of biomass or per individual varies from species to species (Garrido-Sanz, Senar, & Piñol, 2022; Krehenwinkel et al., 2017; Zoschke, Liere, & Börner, 2007), and may also vary among tissues, during development or according to phenology. Two main sources can be considered for technical biases. First, the DNA extraction method, whose efficiency depends on the extracted substrate and varies between taxonomic groups (Dopheide, Xie, Buckley, Drummond, & Newcomb, 2019). Second, the PCR amplification can incur species-specific amplification biases (Pawluczyk et al., 2015) related to the annealing step (Piñol, Mir, Gomez-Polo, & Agustí, 2015) or to the PCR extension step, which may depend, among other things, on the GC content of the metabarcodes (Nichols et al., 2018). Thus, the sum of all these biases obscures the relationship between the abundance of the sequenced reads and the relative abundance of the species in terms of biomass or number of individuals.

Metabarcoding thus requires an appropriate statistical approach to robustly estimate species abundances (Alberdi & Gilbert, 2019; Mächler, Walser, & Altermatt, 2021). For a long time, that quantification problem has been considered. Authors have proposed improvements by optimizing the choice of primers (Krehenwinkel et al., 2017), by varying the number of PCR cycles for different replicates (Silverman et al., 2021) or by creating mock communities to infer correction factors with one species of interest and one control species (Thomas, Deagle, Eveson, Harsch, & Trites, 2016), with two species of interest in different quantities (Matesanz et al., 2019) or by comparing several mock communities of more complex composition (Krehenwinkel et al., 2017); or to infer PCR efficiencies (Shelton et al., 2022). Internal controls can be used, but these do not allow measuring amplification bias (Smets et al., 2016; Ushio et al., 2018).

The present paper examines the biases introduced by the most commonly criticized step of DNA metabarcoding, the PCR amplification. The strength of the amplification bias and its impact on the estimated abundances of metabarcoding are assessed. This study is based on a new mathematical model of PCR amplification that is applicable to the simulation of DNA metabarcoding experiments. Several models exist to describe PCR dynamics (*e.g*. Carr & Moore, 2012; Hayward, 1998; Mehra & Hu, 2005) but have not been linked to metabarcoding. The model developed from existing models considers the amplification bias between species in conjunction with the saturation phase of PCR amplification, with a minimum number of parameters. A usual model in quantitative metabarcoding is the exponential model, also called log-ratio linear model (*e.g*. Gold et al., 2023; Kelly, Shelton, & Gallego, 2019; Shelton et al., 2022), where the abundance of each species increases geometrically during the PCR. The non-treatment of saturation is not a problem in quantitative real-time PCR (qPCR) because the amplification starts with an exponential phase, but is incompatible with metabarcoding PCR, which relies on the final state of the system.

The impact of low priming site conservation on species detection and quantification of COI markers has been widely discussed. These biases are related to the annealing phase of PCR cycles due to primer mismatches (Clarke, Soubrier, Weyrich, & Cooper, 2014; Piñol et al., 2015; Pompanon et al., 2012). To specifically target the biases induced by the extension step of PCR, we assessed them on three mock alpine plant communities using the *Sper01* marker (Taberlet et al., 2007). This marker is widely used in many ecological studies: soil biodiversity (Yoccoz et al., 2012), paleoecology based on ancient eDNA (Willerslev et al., 2014) or diet (Valentini et al., 2009). Although there is very little variation at the *Sper01* priming sites, no strong annealing bias can be assumed for this marker. However, the length of the metabarcodes and the complexity of its sequence (length and frequency of homopolymers) varies from species to species, making it an appropriate candidate to study extension bias. PCR efficiencies for three species were accurately estimated using Taqman qPCR to calibrate our model and then to infer the pre-PCR eDNA proportions of each species. Combined with precise estimates of target DNA concentrations in each species by droplet digital PCR (ddPCR), the results of this experiment demonstrate the benefit of handling PCR extension bias and the variation of target DNA concentration among taxa to correctly estimate taxa abundance from DNA metabarcoding results. Although only a single marker was studied here on a limited number of species, the presented protocol is easily generalizable and opens perspectives for quantitative DNA metabarcoding (qMetabarcoding).

## Material and Methods

### Metabarcoding experiment

Quantification biases were investigated using three mock communities composed of thirteen alpine plants belonging to the *Spermatophyta* clade (Supplementary Table 1), using the *Sper01* primer (Taberlet, Bonin, Zinger, & Coissac, 2018; Taberlet et al., 2007) targeting the P6 loop of the *trn*L of the chloroplast genome. Plant species were selected for having no mismatches at their priming sites with the *Sper01* primers.

### Plant sampling

Plants leaves were collected in Chartreuse and Belledonne massif in the French Alps during Spring 2021 (Supplementary Table 1). Freshly collected material was stored in silica gel before DNA extraction.

### DNA Extraction

Plant DNA was extracted using the CTAB protocol (Doyle, 1990), except for *Carpinus betulus*, for which a *DNeasy Plant Mini Kit* (Qiagen) was used after unsuccessful CTAB extractions.

### Quantification of target DNA

The total DNA concentration for each plant sample was determined using Qubit (ThermoFisher). The amount of DNA targeted by the *Sper01* primer is not proportional to the total DNA concentration, as the number of chloroplasts per cell is expected to vary between different species and tissues and during plant development (Golczyk et al., 2014; Sakamoto & Takami, 2018; Zoschke et al., 2007). ddPCR was used to provide absolute quantification of the *Sper01* target DNA. ddPCR was preferred over qPCR because it is much less affected by inhibition than qPCR, which varies from sample to sample. (Sidstedt, Rådström, & Hedman, 2020). This quantification was performed using serial dilutions of total DNA concentrations ranging from 6.25 × 10^−2^ *ng/μl* to 6.25 × 10^−5^ *ng/μl* with one or two replicates for each condition. The reaction mixtures had a total volume of 20 *μl* (5 *μl* of DNA solution, 10 *μl* of Master Mix EvaGreen, 0.6 *μl* of primers (forward and reverse) at 10*μM*, 4.4 *μl* of milliQ water). The *QX200 Droplet Digital System* (Bio-Rad) was used to generate droplets (*QX200 Droplet Generator*) and to analyze them after PCR amplification (*QX200 Droplet Reader* with the *QuantaSoft Software*). Thermocycler conditions with optimized annealing temperature for the *Sper01* primer (52^*°*^C) were set (30 seconds at 95^*°*^C, 30 seconds at 52^*°*^C, one minute at 72^*°*^C). Replicates identified as incorrect by the reader and the most diluted replicate in cases where this concentration was outside the expected detection range were removed.

In this study, the concentration index chosen to compare the samples is the expected number of target copies per *ng* of total DNA. It is calculated from each assay as in the equation 1. The number of copies per *μ*l (in target DNA) is the value measured by ddPCR. C(Total DNA)_*replicate*_ is the total DNA concentration of the sample in the reaction mix. The average concentration for each species is used for the rest of the protocol.

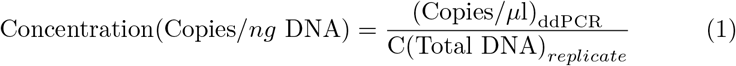

This choice of index was made because mock communities are composed of purified plant DNA extracts. This correction includes two bias factors: the number of target copies per genome and the genome size. In the case of an eDNA sample, the amount of target DNA detected does not depend on the genome size. The aim is to quantify the number of sampled cells, so the only factor to correct is the difference in copy numbers per genome. Two choices are then possible. Either, the concentration is established in copies per gram of tissue (equation 2), and then (Copies/*μ*l)_ddPCR_ is calculated for a known mass of tissue m(Tissue)_*replicate*_. Or the concentration is established by taking into account the genome size using the C-value (equation 3, Supplementary Table 1) with the same ddPCR assays as in equation 1. The Kew C-value database (https://cvalues.science.kew.org/) can be used to implement this correction, possibly using a parent species as an approximation.

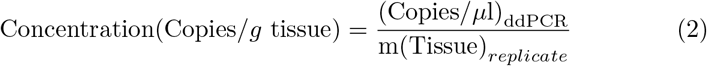

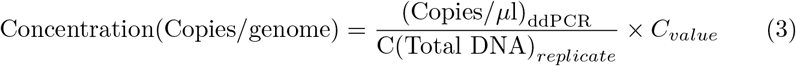

### Mock communities

Three mock communities were constructed after the ddPCR assays: (i) a uniform community (**ℳ**_***U***_) where each plant has the same concentration of target DNA, (ii) a community where each plant has the same concentration of total DNA (**ℳ**_***T***_), and (iii) a community where the concentrations of target DNA are distributed according to a geometric sequence of common ratio 1/2 (concentrations of 1, 1/2, 1/4…) (**ℳ**_***G***_). The species used are described in Table 1. The metabarcode sequences are given in the Supplementary Table 1 and the exact composition of each community is given in the Supplementary Table 2. The comparison between **ℳ**_***U***_ and **ℳ**_***T***_ communities allows determining the bias introduced by variation in the number of chloroplast genomes per unit of total DNA. The **ℳ**_***U***_ and **ℳ**_***G***_ comparison allows the estimation of relative PCR extension step efficiencies.

**Table 1:**
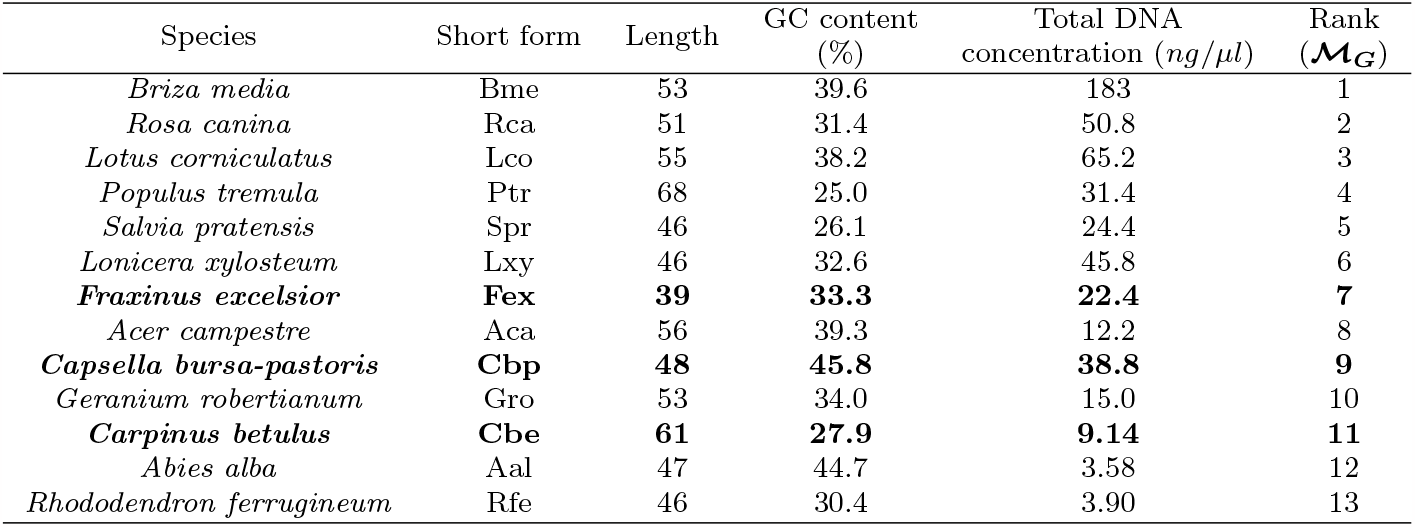
Plants used for the three mock communities and their characteristics for the *Sper01* marker. Total DNA concentrations are assayed in the samples after extraction by Qubit. Rank stands for decreasing abundance in the **ℳ**_***G***_ community.

**Table 2:** Proportions in **ℳ**_***U***_ and relative PCR amplification efficiencies measured for the four Taqman qPCR probes and inferred from the **ℳ**_***U***_ community. The maximum efficiency was set at 1 for *Rosa canina*. Efficiencies inferred were normalized so that Fex has the same efficiencies with both methods.

| Species | Average proportion in $\mathcal{M}_U$ (%) | | PCR Efficiency inferred from | |
| --- | --- | --- | --- | --- |
| | Theoretical | Observed | Taqman | $\mathcal{M}_U$ |
| Bme | 7.7 | 6.1 |  | 0.918 |
| Rca | 7.7 | 17 |  | 1.00 |
| Lco | 7.7 | 8.0 |  | 0.939 |
| Ptr | 7.7 | 8.6 |  | 0.945 |
| Spr | 7.7 | 2.7 |  | 0.855 |
| Lxy | 7.7 | 5.5 |  | 0.910 |
| **Fex** | **7.7** | **6.3** | **0.920** | **0.920** |
| Aca | 7.7 | 8.3 |  | 0.942 |
| **Cbp** | **7.7** | **11** | **0.968** | **0.961** |
| Gro | 7.7 | 2.4 |  | 0.847 |
| **Cbe** | **7.7** | **8.1** | **0.951 (CbeA)**<br>**0.927 (CbeB)** | **0.940** |
| Aal | 7.7 | 5.8 |  | 0.914 |
| Rfe | 7.7 | 10 |  | 0.958 |

### DNA metabarcoding PCR amplification

For each community, 20 replicates (2*μl* of DNA) and one PCR negative control (2*μl* of milliQ water) are made. Three wells are left blank (sequencing controls). Each well was individually tagged. 40 PCR cycles were run with an optimized annealing temperature for *Sper01* (30 seconds at 95^*°*^C, 30 seconds at 52^*°*^C, one minute at 72^*°*^C).

### Metabarcoding DNA Sequencing

High-throughput sequencing was performed on NextSeq (Illumina) by Fasteris (Plan-les-Ouates, Switzerland; https://www.fasteris.com/). One library was constructed per community following the Metafast protocol (as proposed by Fasteris).

### Bioinformatic pipeline

All the bioinformatic work was performed on a laptop MacBook Air (2017, 2.2 GHz Intel Core i7 Dual Core Processor). The data and analysis scripts are available on the project’s git page, https://github.com/LECA-MALBIO/metabar-bias. Raw data was processed with OBITools (version 4 aka OBITools4; Boyer et al., 2016, https://metabarcoding.org/obitools4). Unless otherwise stated, the further analyses were carried out using R.

### A DNA metabarcoding experiment model

The goal of this part is to estimate the initial relative abundances *p*_*s*_ of each species *s*, from the number of reads *R*_*s*_ among the *S* different species in the considered environmental sample. To achieve this, a simulation model was used to generate virtual metabarcoding data that are compared with observed data.

The model integrates the three steps involved in the production of a DNA metabarcoding result from a DNA extract, as in Gold et al. (2023): i) the sampling of a portion of the DNA extract, ii) the PCR amplification, iii) the sampling of a portion of the PCR reaction for sequencing.

### Sampling of a portion of the DNA extract

The initial number of molecules in a replicate 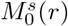, is modeled by a Poisson distribution with expectation 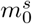.

On the basis of simulations, it was observed that the final observed proportions had a standard deviation about 25 times higher than the proportions in equivalent simulated data. Such a standard deviation can be obtained in simulations by replacing the Poisson distribution with a negative binomial distribution with parameters 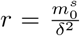 and 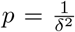, so that its expectation is 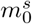 and its variance is 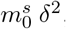, with *δ* the overdispersion of the standard deviation obtained with an initial Poisson distribution (with here *δ* ≃ 25). Our choice for the Poisson distribution simplifies the model and the mean value remains unchanged.

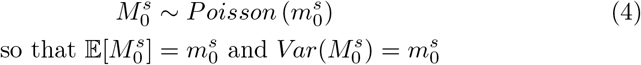

### PCR amplification

The used PCR model, here called logistic model, accounts for the different amplification efficiencies and the saturation phase. The model is an adaptation of the deterministic model in Hayward (1998), which also features a logistic form with an additional parameter. It is also a simplified form of the model in Carr and Moore (2012). Compared with a conventional exponential model, the logistic model accounts for saturation phase at the end of the PCR. Both are parametric stochastic models. Figure 1 illustrates kinetics described by these two parametric stochastic models. The models are fitted to qPCR data generated from a sample of *Capsella bursa-pastoris*. The fit is a non-linear regression with more weight given to the acceleration cycles (15^th^-25^th^ cycles).

**Fig. 1:**
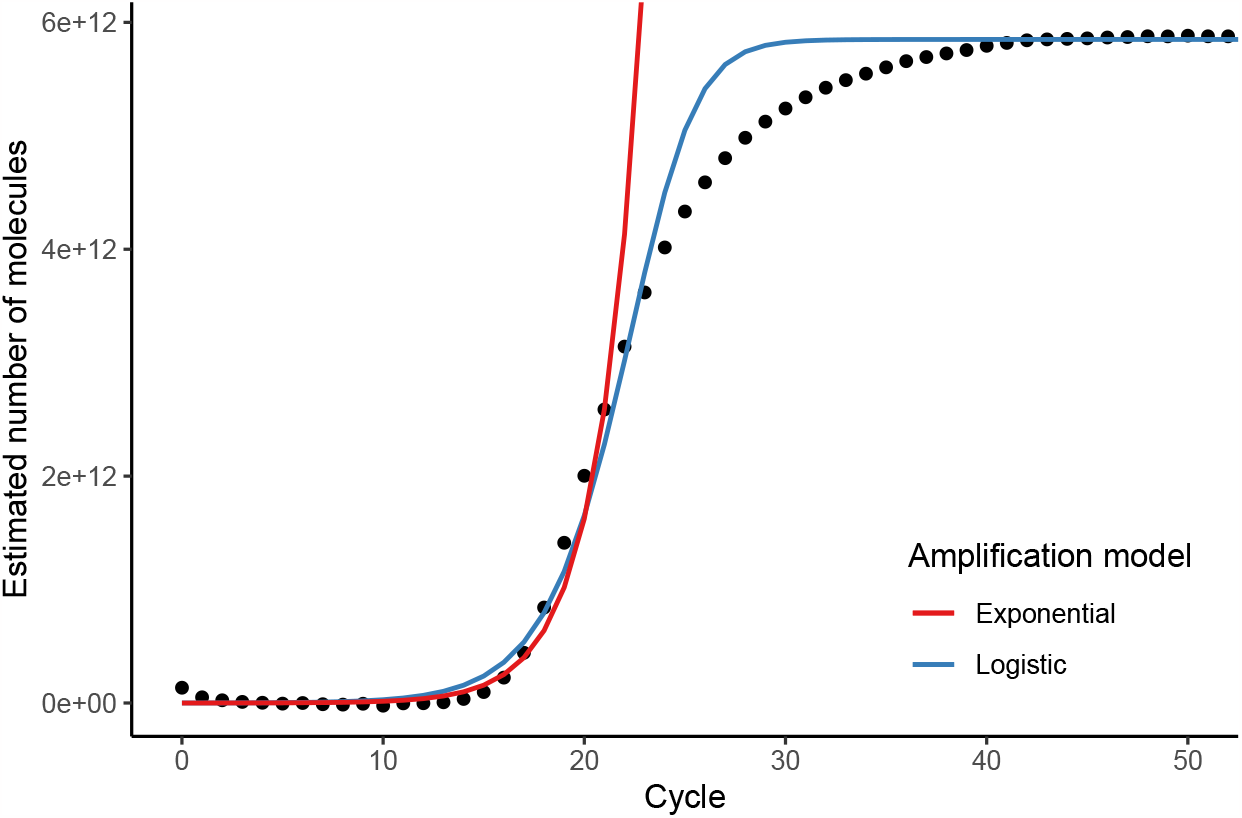
Observed qPCR kinetics for a sample of *Capsella bursa-pastoris* (black dots) compared to two PCR models fitted to the data. Blue curve: logistic model; red curve: exponential model. An asymmetry of amplification is observed around the inflection point, which creates here a gap between the 25^*th*^ cycle and the 40^*th*^ cycle for the logistic model. This asymmetry is taken into account in more sophisticated PCR models (Gottschalk & Dunn, 2005). It is known that the first qPCR cycles correspond to a RFU background noise (Rao et al., 2013).

The models considered describe the evolution of the number of DNA molecules of each species cycle by cycle, denoted 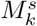for each species *s* at PCR cycle *k*. Each molecule already present is maintained and has a probability 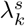 of being replicated again, modeled by a binomial distribution (equation 5) depending on the state of the system after cycle *k* − 1. More precisely, amplification of species *s* depends on 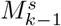 and on 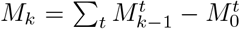 because the saturation depends on all the molecules that were created before cycle *k* (equation 6).

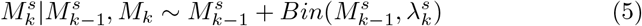

Let *K* be a charge capacity *K, ie* the total number of DNA molecules that can be created during the amplification. This quantity is not known a priori. Due to saturation, the effective PCR efficiency of each species, 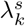, decreases during the PCR. The logistic saturation has been chosen for its simple shape (equation 6).

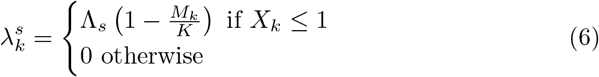

The purely exponential model is a special case with no saturation where 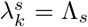 at each cycle *k*. Under the exponential model, the expected number of molecules at cycle *k* is given by equation 7.

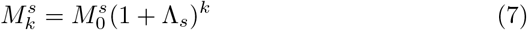

### Sampling of a portion of the PCR reaction for sequencing

All the molecules created by the PCR are not sequenced: only a fraction constitutes the observed data, denoted *R*_*s*_ for each species *s*. At the end of a PCR amplification with *n* cycles, the sequencing step is described as a multinomial sub-sampling of *R*_total_ molecules (equation 8).

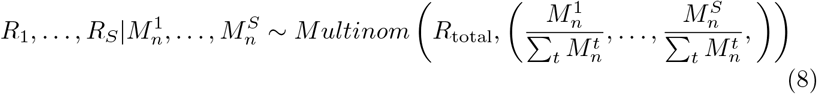

### Measure of the amplification efficiencies Using Taqman qPCR assay

PCR amplification efficiencies Λ_*s*_ were measured by qPCR for three of the plant species present in our mock communities: *Carpinus betulus, Capsella bursa-pastoris* and *Fraxinus excelsior*. These three species were chosen because their metabarcodes differ widely in sequence length and GC content. This makes it possible to expect different amplification efficiencies and to design specific Taqman internal probes that allow individual PCR efficiency measurements within a mixture of the three plant DNAs. Two different probes were designed for *Carpinus betulus* to evaluate the influence of the probe itself on the measurement. The four probes used are described in the Supplementary Table 3. The assay was performed using Taqman qPCR on a uniform community composed of these three species. A 5-fold serial dilution from 654 to 1 copies/*μl* in the reaction mix (25 *μl* with 5*μl* of DNA) was performed for each probe, with three replicates per concentration. Taqman qPCR was chosen to measure PCR efficiency because it allows measurement from a mixture of the three plant DNAs. This ensures the same inhibitory effect for each species. Since each individual DNA extract has its own pool of inhibitors that interfere with qPCR assays, independent measurement on pure extract would not be realistic (Svec, Tichopad, Novosadova, Pfaffl, & Kubista, 2015). This approach uses the quantification cycles *C*_*t*_ measured by qPCR.

**Table 3:** Proportions of species in **ℳ**_***T***_ and **ℳ**_***G***_. Inferred with **ℳ**_***U***_ means corrected by the ratios. Proportions inferred with Λ_*s*_ are obtained by fitting the PCR model using the efficiencies inferred previously.

| Species | Average proportion in $\mathcal{M}_T$ (%) | | | | Average proportion in $\mathcal{M}_G$ (%) | | | |
| --- | --- | --- | --- | --- | --- | --- | --- | --- |
| | Theoretical | Observed | Inferred with $\mathcal{M}_U$ | Inferred with $\Lambda_s$ | Theoretical | Observed | Inferred with $\mathcal{M}_U$ | Inferred with $\Lambda_s$ |
| Bme | 8.5 | 6.2 | 8.4 | 8.3 | 50 | 36 | 54 | 53 |
| Rca | 14 | 26 | 12 | 12 | 25 | 40 | 20 | 20 |
| Lco | 9.5 | 11 | 11 | 12 | 13 | 15 | 16 | 16 |
| Ptr | 17 | 21 | 20 | 20 | 6.3 | 5.2 | 5.5 | 5.6 |
| Spr | 10 | 3.9 | 12 | 11 | 3.1 | 0.63 | 2.0 | 2.1 |
| Lxy | 5.0 | 2.7 | 3.8 | 3.9 | 1.6 | 0.94 | 1.5 | 1.5 |
| **Fex** | **5.9** | **4.3** | **5.7** | **5.7** | **0.78** | **0.68** | **0.97** | **0.95** |
| Aca | 9.1 | 7.9 | 7.8 | 8.2 | 0.39 | 0.16 | 0.17 | 0.17 |
| **Cbp** | **2.6** | **3.2** | **2.4** | **2.5** | **0.20** | **0.19** | **0.15** | **0.16** |
| Gro | 5.6 | 2.1 | 6.5 | 7.1 | 0.098 | 0.019 | 0.064 | 0.085 |
| **Cbe** | **3.6** | **2.8** | **2.6** | **2.7** | **0.049** | **0.030** | **0.031** | **0.034** |
| Aal | 6.1 | 1.5 | 2.3 | 2.1 | 0.024 | 0.0045 | 0.0072 | 0.0049 |
| Rfe | 2.5 | 5.1 | 3.7 | 3.9 | 0.012 | 0.014 | 0.012 | 0.014 |
| AbsErr |  | 0.045 | 0.017 | 0.018 |  | 0.057 | 0.020 | 0.019 |
| RelErr |  | 0.53 | 0.26 | 0.28 |  | 0.50 | 0.34 | 0.33 |
| $^1D$ | 11 | 8.9 | 11 | 11 | 4.0 | 3.7 | 3.7 | 3.8 |
| $^2D$ | 10 | 6.7 | 9.1 | 9.1 | 3.0 | 3.1 | 2.8 | 2.8 |

The exponential model (equation 7), which is valid before the PCR saturation phase, can be used to estimate apparent PCR efficiencies. Estimated efficiencies are referred to as apparent efficiencies because inhibition is always present. For this study, however, only the relative values of the efficiencies are important. A commonly used formula (equation 9, Gill, Bleka, & Fonneløp, 2022) can be derived from the exponential model to estimate amplification efficiencies from a series of qPCRs performed on successive dilutions. However, a major limitation of this formula that has been identified here is that the estimation of the slope is very sensitive to small variations in *C*_*t*_, resulting in a large variance of the estimator of the efficiency Λ.

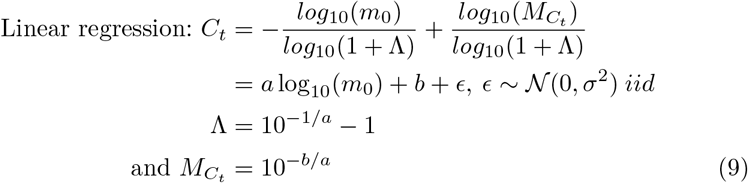

To estimate efficiencies more precisely, this approach is modified. This linear regression approach was used only to estimate 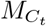, the number of molecules present at *C*_*t*_. This quantity was assumed to be shared by all species. The value retained was the average of the estimated value for each species: 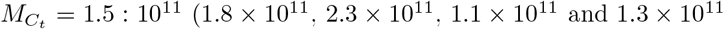 and 1.3 × 10^11^ for probes CbeA, CbeB, Cbp and Fex, resp.) (equation 9).

From this value of 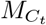, other quantities can be estimated: *K*, the total number of molecules that can be created during amplification, and Λ_*s*_, the PCR efficiencies.

The estimate of *K* is given by equation 10. This equation is established from observed relative fluorescence unit (RFU) values, assuming within-replicate proportionality between RFU and DNA copy number (Gill et al., 2022), although RFU values are not standardized, and fluorescence saturates at the end of amplification and depends on many experimental factors (Svec et al., 2015). This approach only provides an estimate: *K* = 7.6 × 10^12^ here (see Discussion). For the inference of initial abundances, the value of *K* used was estimated numerically (see below).

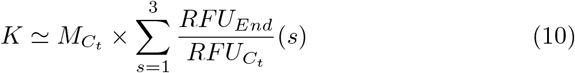

Then, the efficiencies Λ_*s*_ were estimated for each replicate, for which the initial quantities 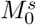 are known (equation 11). For subsequent analyses, the average Λ_*s*_ over all replicates is used.

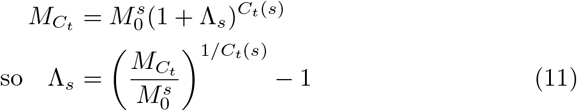

If the value of 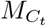chosen is not the average but the minimum value 1.1 × 10^11^ (resp. maximum value 2.3 × 10^11^), the inferred values of Λ_*s*_ are multiplied by a factor of 0.980 (resp. 1.03).

### Using the ℳ_*U*_ community

PCR efficiencies Λ_*s*_ were also inferred by fitting the logistic PCR model presented above to the experimental data. *K* is also inferred at the same time in this approach. For the **ℳ**_***U***_ community, the known quantities are the read numbers *R*_*s*_ and the initial quantities 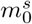 of each species. The Fixed Landscape Inference MethOd (*flimo*, Moinard, Oudet, Piau, Coissac, & Gonindard-Melodelima, 2022) implemented in Julia was used for this purpose. This algorithm is based on model simulations compared with observed data using summary statistics, in the same way as ABC methods. It has the advantage of being faster. The *flimo* method minimizes an objective function in the form of a *χ*^2^ statistic (equation 12).

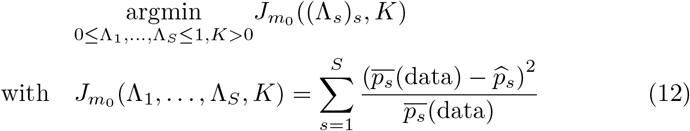

where 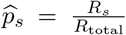 is the average proportion of species *s* in a replicate, estimated over *n*_*sim*_ = 190 simulations for given 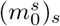, (Λ_*s*_)_*s*_ and *K*, and *p*_*s*_(data) is the average proportion of species *s* in the data.

The inferred efficiencies are relative, as the model can produce similar results for different ranges of Λ_*s*_, especially when the value of *K* changes. The maximum efficiency value has been set at 1. 101 parameter inferences were performed. For the inference of initial proportions in **ℳ**_***T***_ and **ℳ**_***G***_, the values retained for Λ_*s*_ and *K* are those associated with the median value of *K*.

### Correction of relative abundances of a MOTU

Figure 2 summarizes the additional pipeline recommended for correcting amplification bias in a metabarcoding experiment. The PCR amplification efficiency of each species is estimated from samples of species characteristic of the ecosystem studied that are assayed by ddPCR. There are two ways of doing this: Taqman qPCR or a mock community study. These efficiencies are then used to infer the initial proportions of each species.

**Fig. 2:**
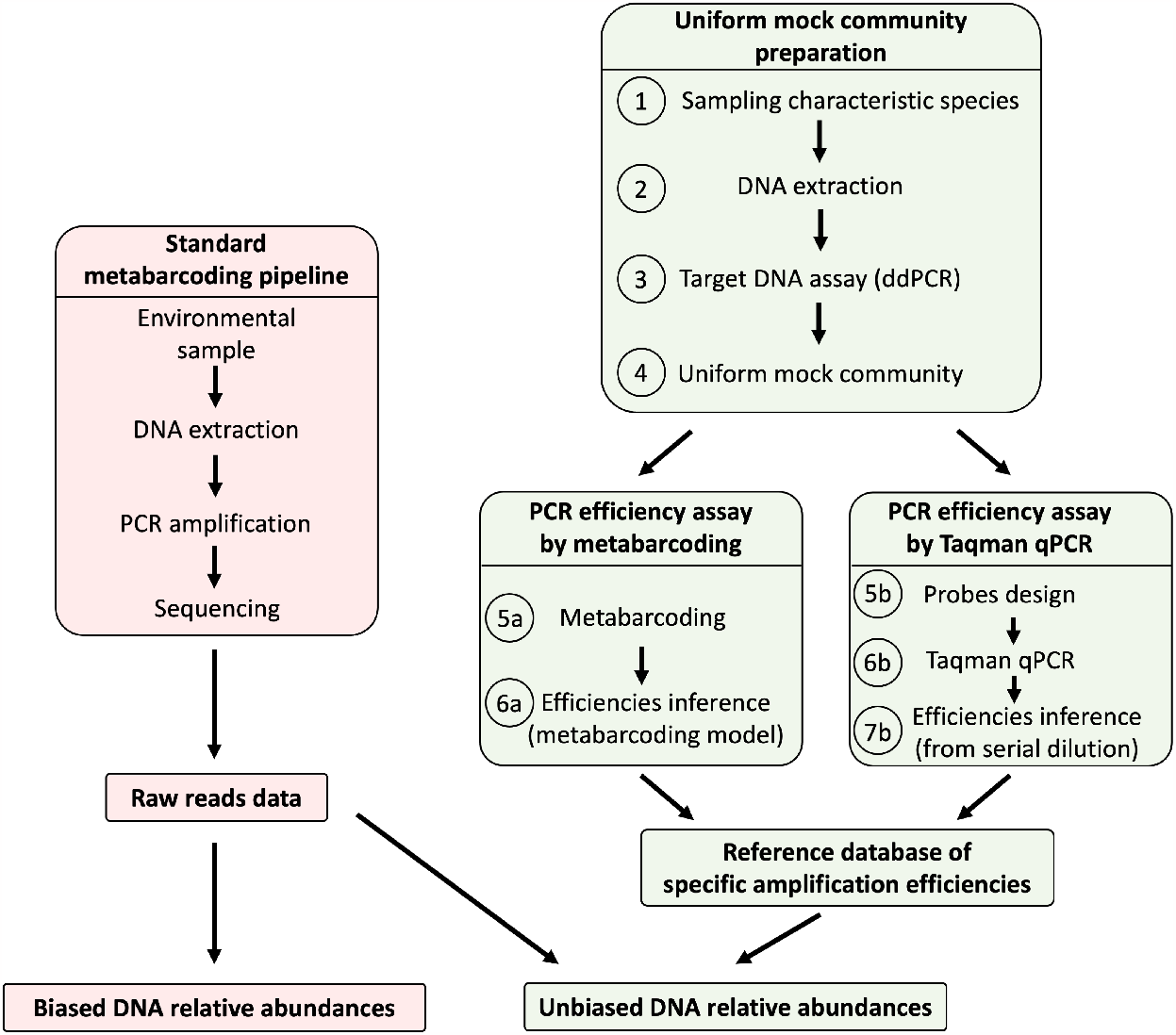
Additional pipeline recommended for correcting amplification bias in a metabarcoding experiment as presented in this study.

### Using the Ratio method

Previous works (*e.g*. Shelton et al., 2022; Silverman et al., 2021) showed that a reference mock community can be used to correct abundances in another community composed of the same species. Although this was not the main objective of our work, this result was verified using the three communities studied. The **ℳ**_***U***_ community was used as a reference to correct abundances in the **ℳ**_***T***_ and **ℳ**_***G***_ communities. In the **ℳ**_***U***_ community, each species had a starting relative frequency of 1*/*13 ≃ 7.7%, which should have been observed in the final read proportions in the absence of amplification bias. The correction factor for each species *c*_*s*_ is therefore simply the median ratio between the expected and the observed reads frequencies over all replicates in the **ℳ**_***U***_ community (equation 13).

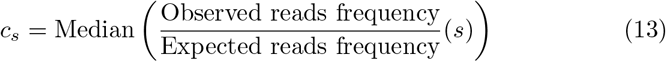

For the **ℳ**_***T***_ and **ℳ**_***G***_ communities, this correction factor is applied to estimate the initial proportions 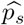 for each species *s* with *R*_*s*_ reads (equation 14).

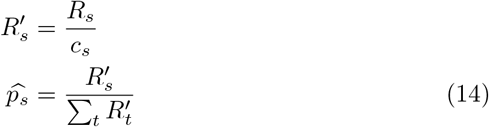

### Using the estimated amplification efficiencies

The inference of the actual proportions of eDNA from the relative read abundances (RRA) measured after DNA metabarcoding sequencing is achieved by the same algorithmic method presented above, but this time the Λ_*s*_ efficiencies are assumed to be known, either measured by Taqman qPCR or inferred from the model fit for the **ℳ**_***U***_ community. The parameters to be inferred are the initial quantities 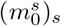 for the **ℳ**_***T***_ and **ℳ**_***G***_ communities (equation 15).

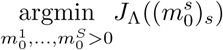

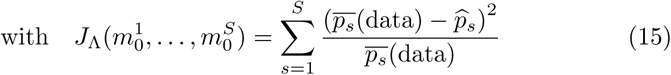

An estimate of these proportions can be obtained using the exponential model, but this requires knowledge of the PCR effective number of “exponential cycles” at saturation (equation 17) which cannot be inferred simultaneously to Λ_*s*_ and *K* with the exponential model. In average, the final proportion *p*_*s*_ of each species *s* is:

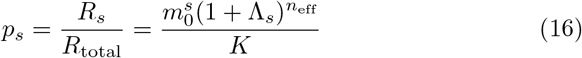

Taking the logarithm and summing all the species :

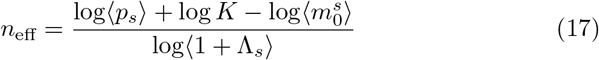

where, 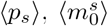 and ⟨1 + Λ_*s*_⟩ are the geometric mean of the 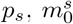 and 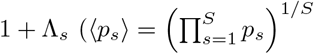, etc.).

### Criteria for measuring quantification errors

The distance between the observed or corrected proportions 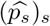, median over all the replicates) and the initial theoretical proportions 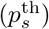 is measured by two RMSE (*Root-Mean-Square Error*) criteria. The error measured is either absolute (equation 18) or relative (normalized by the theoretical proportions, equation 19).

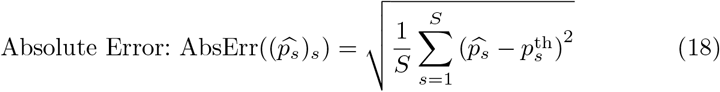

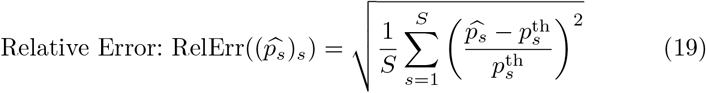

### Ecological conclusions: biodiversity indices

To compare theoretical, observed and inferred compositions, biodiversity indices were computed for **ℳ**_***T***_ and **ℳ**_***G***_. Hill numbers (Hill, 1973) (equation 20), interpretable as an effective number of species in the community, were chosen with *q* = 1 (linked to Shannon entropy) and *q* = 2 (linked to Gini-Simpson index).

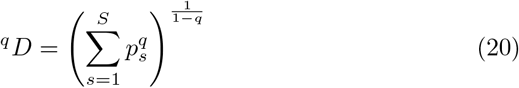

## Results

### ddPCR assay

The concentrations of each plant sample measured by ddPCR are shown in Figure 3. For the same total DNA concentration, there was a wide variability in average target concentration, ranging from 3.7 × 10^4^ copies per *ng* for *Rhododendron ferrugineum* to 2.5×10^5^ copies per *ng* for *Populus tremula* with an average of 1.1 × 10^5^ copies per *ng* among the thirteen species. The factor between the extremes is thus 6.6.

**Fig. 3:**
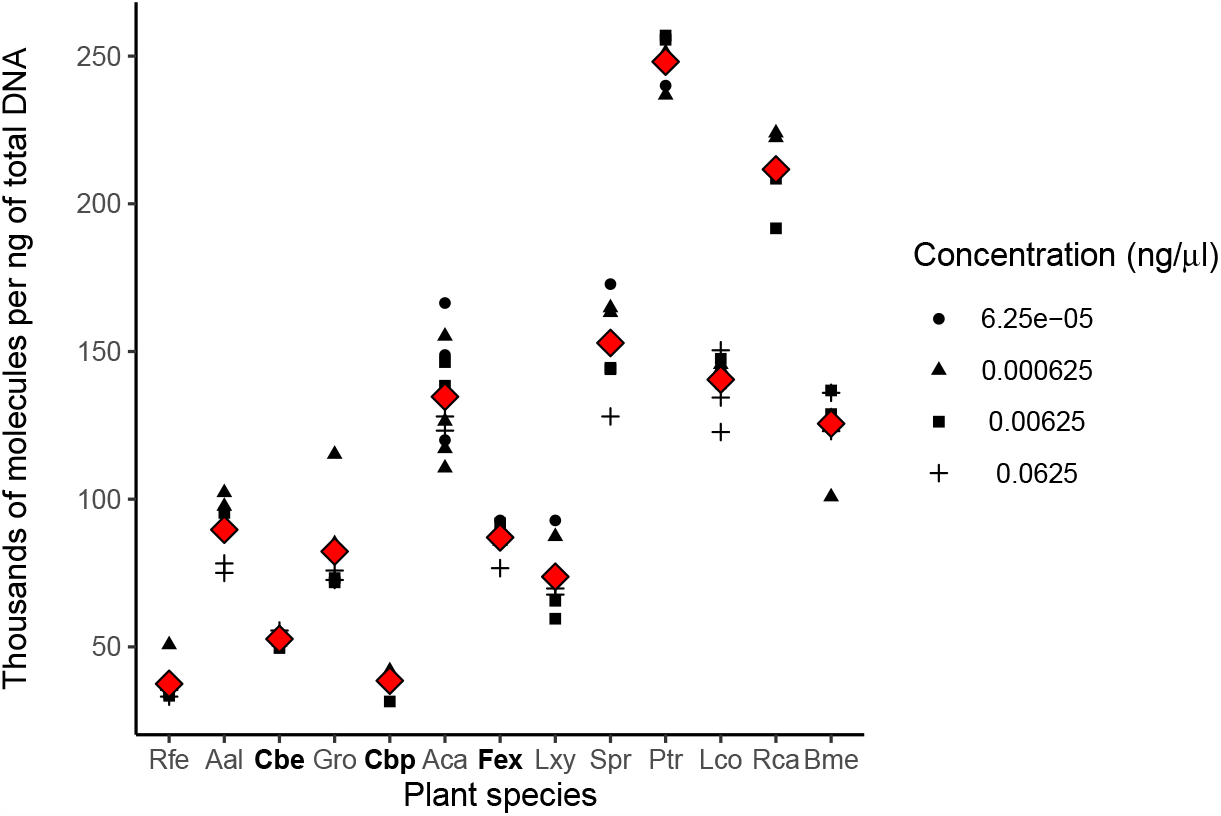
Number of target DNA molecules (thousands) per *ng* of total DNA for thirteen alpine plants, computed with the index used in equation 1. Each black dot is a replicate, for different total DNA concentrations. The red diamonds correspond to the mean for each species.

### Metabarcoding experiment

#### Raw sequencing data

After processing with the OBITools, an average of 37,000 reads per non-negative replicate was obtained with a standard deviation of 27,000 reads (first and third quartiles : 14,000 and 56,000 reads). Negative controls showed negligible contamination. For each community out of the 20 PCR replicates, one replicate with fewer than 5,000 reads was discarded from further analysis.

#### Reads proportions

The comparison of observed and expected read proportions is shown in Figure 4. Significant differences can be observed: at most, between the observed and expected proportions, there is a factor of 3.0 for *Geranium robertianum* in the **ℳ**_***U***_ community, 4.2 for *Abies alba* in **ℳ**_***T***_ and 9.0 for *Abies alba* in **ℳ**_***G***_.

**Fig. 4:**
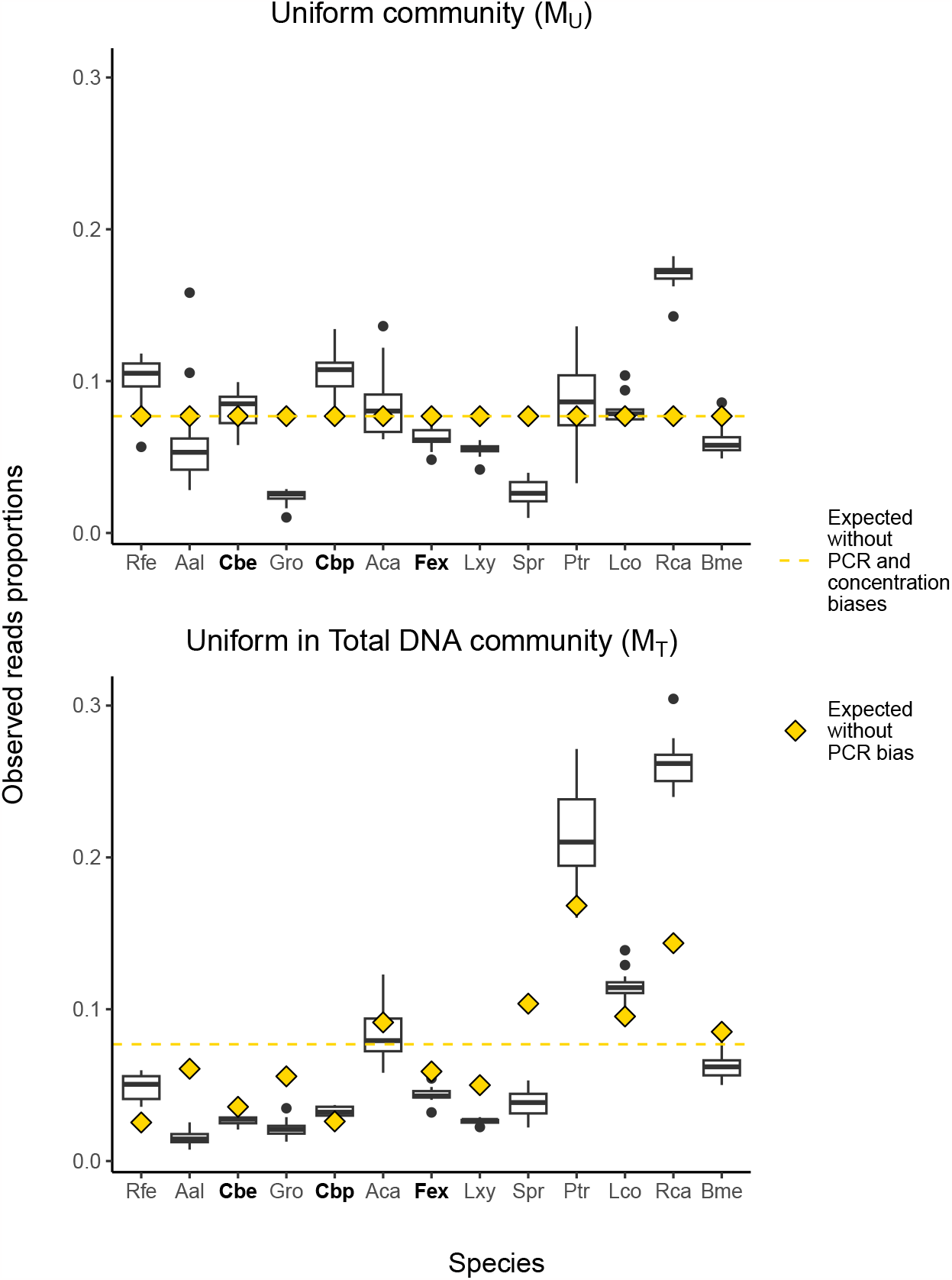
Observed relative proportions of reads of thirteen plant species for the mock communities **ℳ**_***U***_ and **ℳ**_***T***_. Gold lines indicate proportions expected in the absence of target concentration and amplification bias. Gold diamonds are the proportions expected in the absence of amplification bias. For the **ℳ**_***U***_ community, the deviation of the boxplots from the diamonds shows the amplification bias alone. For the **ℳ**_***T***_ community, both biases are present. Concentration bias is visible as the difference between the diamond and the line.

Comparing the observed proportions with the expected proportions allows to visualize the two biases under study. For example, *Rosa canina* species has both good efficiency and a high target concentration: the two biases add up. Conversely, *Geranium robertianum* is penalized by both biases. *Salvia pratensis* has a higher-than-average concentration, but poor efficiency. *Capsella bursa-pastoris* is well amplified, but its target concentration is low.

The joint effect of the double bias is visible for **ℳ**_***T***_, with median proportions comprised between 1.5% and 26%, and between 2.6% and 17% for **ℳ**_***U***_.

Inter-replicate variability is significant in some species, such as *Populus tremula* (in **ℳ**_***U***_ : mean proportion : 8.6%, varying from 3.3% to 14%, standard deviation of 2.7%).

### Inferring PCR efficiencies and abundances

The apparent PCR efficiencies for the three species tested (**Fex, Cbe, Cbp**) measured using the Taqman qPCR method for the four probes have a relative difference of the order of 5%. That can be considered low, but due to the exponential nature of PCR, it has a real impact on the final proportions in the community due to the exponential nature of PCR amplification.

Table 2 shows the abundances in the reference mock community **ℳ**_***U***_ and the efficiencies inferred from the Taqman qPCR assay and from the model fit to the **ℳ**_***U***_ community, with the *flimo* method. The standard deviation of the PCR efficiencies established for each probe by Taqman qPCR varies between 0.0049 (Cbp) and 0.011 (Fex). When inferred from **ℳ**_***U***_, the standard deviation of Λ_*s*_ is between 0.0027 (Cbp) and 0.010 (Gro). The lowest efficiency is around 15% lower than the maximum. The absolute values determined by Taqman qPCR are overestimated in relation to these values, but once normalized, they are broadly similar, even though more values would be required for a rigorous comparison. Because of this similarity and the fact that the assay involves only three species, the results are based on efficiencies measured in **ℳ**_***U***_.

The inferred median K value is 2.36 × 10^12^ (s.d. = 2.61 × 10^12^). The choice of the value of *K* has a negligible influence on the inferred values of Λ_*s*_.

Table 3 shows the proportions in the **ℳ**_***T***_ and **ℳ**_***G***_ communities, as well as the errors compared to the theoretical proportions and the biodiversity indices.

The standard deviation of inferred proportions varies from 0.28% (Lco) to 2.0% (Gro) of the mean inferred proportion for **ℳ**_***T***_ (resp. from 0.11% (Lco) to 7.0% (Aal) of the mean inferred proportion for **ℳ**_***G***_). The results of the two corrections are comparable and both improve the RMSE criteria, as expected. The corrected biodiversity indices also seem to better approximate the real biodiversity than the observed values.

### PCR bias importance: comparison of model simulations and observed data

To illustrate the effect of small differences in efficiency, PCR kinetics were simulated for two species with equal initial quantities. Figure 5 shows the final proportions of the two species according to the difference in PCR efficiency. These simulations are compared with the proportions observed in the **ℳ**_***U***_ community when comparing *Rosa canina* (the most efficiently amplified species) and the other species individually. These two proportion series are very close to each other.

**Fig. 5:**
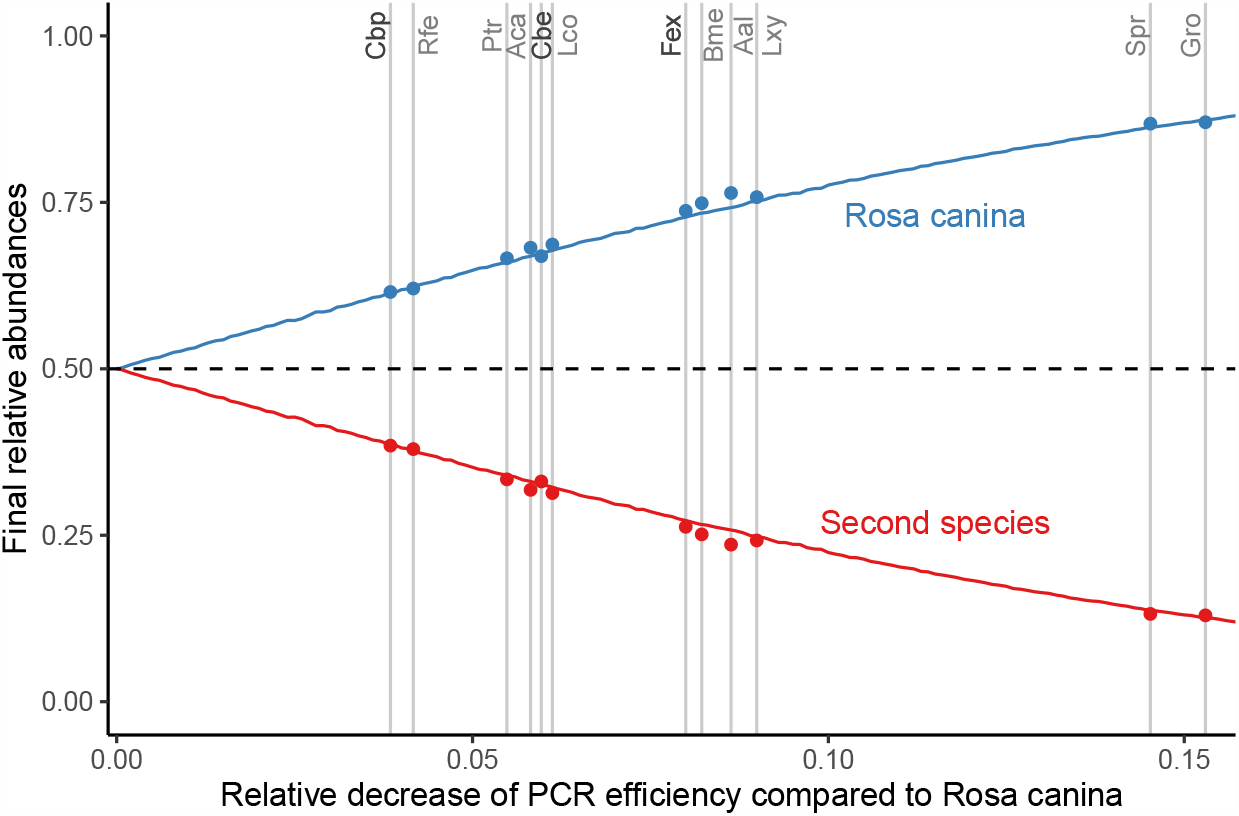
Relative abundances in a mock community of two initially evenly distributed species simulated with the logistic model (lines) and observed in the **ℳ**_***U***_ community (dots) considering only *Rosa canina* and the other species individually. The first species has an efficiency of Λ_1_ = 1. The second has a variable efficiency, of value Λ_2_ = Λ_1_(1 − *x*) along the *x*-axis (Λ_2_ ∈ [0.85, 1.0]).

## Discussion

The quantitative aspect of DNA metabarcoding is regularly questioned by ecologists. Here, two potential biases were considered and their relative effects quantified.

The first is well known. It has long been discussed by microbial ecologists (Kembel, Wu, Eisen, & Green, 2012; Milivojević et al., 2021) and has been identified for macro-organisms (Garrido-Sanz et al., 2022; Krehenwinkel et al., 2017). It can be summarized by a simple question: how many copies of the target gene marker are present per genome in each species under consideration?

In macro-organisms such as plants and animals, most of the targeted markers are carried by the chloroplast or mitochondrial genome, but the same question remains: how many copies of the organelle genome are there per cell? This amount can be estimated by ddPCR. In this study, the communities studied were composed of purified DNA, assayed by ddPCR in marker copies per *ng* of total DNA. In the case of an environmental DNA sample, this measurement is easily adapted by taking into account the C-value (equation 3) to assay the target in marker copies per genome. Another option is to measure the target in copies per unit of biomass, as in Thomas et al. (2016) or (Matesanz et al., 2019) (equation 2) Such a reference database makes it possible to convert molecule proportions into species biomass proportions. Here is an example of the bias in marker copies per genome. Among the 13 plants tested, the one more concentrated in chloroplast DNA, *Populus tremula* (Ptr), has 6.6 times more copies per unit of nuclear DNA than the one less concentrated, *Rhododendron ferrugineum* (Rfe). According to the Kew C-value database, the 1C value of Ptr is 0.45 pg (Siljak-Yakovlev et al., 2010) and that of *Rhododendron ponticum*, the only *Rhododendron* measured, is 0.74 pg (Bou Dagher-Kharrat et al., 2013) (See Supplementary Table 1). Both together allow to estimate that the bias in chloroplast abundance (in copies per genome) can lead to a 4-fold overestimation of Ptr abundances relative to Rfe (equation 2).

The second type of bias is amplification bias, which our study quantifies precisely. The amplification efficiency of a marker for the species *s* (Λ_*s*_) is an intrinsic property of the sequence. It does not depend on co-amplified sequences. In this study, we propose two methods to measure it. Both provide similar values, and the choice between them depends on practical convenience. The values obtained can be used to correct the composition of any community, as long as differences in amplifiability between the species present do not cause one or more to disappear. The proposed correction method combines the generation of a reference base for the amplifiability and a mathematical model of the PCR. It does not require any modification of the metabarcoding protocol. Therefore, it can be applied to already generated results and is easy to implement. In particular, experimental parameters do not need to be adjusted to keep amplification within the exponential regime. The logistic model is valid even if saturation is not reached, *i.e*. if 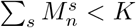*K*. However, it is necessary to achieve the saturation plateau while amplifying the reference community (**ℳ**_***U***_ in this case) in order to infer the efficiencies of Λ_*s*_ and *K*. Without obtaining the saturation plateau, one cannot determine the stage at which the PCR was terminated. Estimating *K* from Taqman qPCR data is imprecise, and numerical inference along with Λ_*s*_ seems more appropriate. The values of Λ_*s*_ are relative and vary with *K* linearly: this is why a precise value of *K* is not necessary to establish a correction.. This second method suggests that a precise *K* value is not necessary to determine the Λ_*s*_. The order of magnitude is the same for both approaches and is comparable to the estimate of Newton and Graham (2000) (1.81 × 10^12^ molecules at the end of the PCR).

The amplification bias is accumulated over each PCR cycle. Thus, the final bias on the observed read relative frequencies is a function of the amplifiability per cycle and the number of amplification cycles. In PCR, the actual number of amplification cycles is not necessarily the number of cycles programmed into the PCR instrument. This number may be lower because the total amount of DNA that can be synthesized is limited by the nucleotide concentration. It is therefore possible that the plateau will be reached before the programmed number of cycles has been reached, with the last cycles not corresponding to any amplification (Figure 1). Correcting for bias using the ratio method (*e.g*. Shelton et al., 2022; Silverman et al., 2021) requires that each sample, including the reference mock community used to estimate it, be amplified with the same effective number of PCR cycles *n*_eff_. For the exponential model, which is an assumption of the ratio method (Luo et al., 2022), the change in *n*_eff_ can lead to significant variations in the inferred proportions. This means that each sample must contain the same total number of target DNA molecules at the start of the PCR. In our study, each mock community was prepared with close total amounts of target DNA, thus respecting the ideal condition for using the ratio method. Therefore, as shown in Table 3, the corrections made by the ratio method and our PCR model-based approach are strictly equivalent. When samples contain different amounts of target DNA, the efficiency of the ratio method decreases because the number of effective PCR cycles varies from sample to sample. To illustrate, using our logistic PCR models, DNA metabarcoding experiments were simulated for two uniform communities containing 10 times less or 10 times more initial DNA molecules than **ℳ**_***U***_. The maximum final ratio observed from simulation (Rca to Gro) is 9.5 in the former case (abundances of 19% versus 2.0%) versus 5.4 in the latter (abundances of 16% versus 2.8%), while it was 6.7 in the original **ℳ**_***U***_ (abundances of 17% versus 2.6%). Fortunately, oppositely to ratio method, our PCR model-based correction allows us to estimate the effective number of PCR cycles for each sample, thereby accounting for sample heterogeneity, making our correction method more robust.

When the two species with the most different amplifiability, *Rosa canina* (Λ_*Rca*_ = 1.000) and *Geranium robertianum* (Λ_*Gro*_ = 0.847) are co-amplified, with equal amounts of initial target DNA in the extract, the ratio between the RRA observed after sequencing can be up to 6.7 (Figure 5), leading to a strong overestimation of Rca abundance relative to Gro. This initial assessment shows that due to the exponential nature of PCR, even a small difference in amplifiability, as little as 15% between Rca and Gro, the two extreme species tested, can have as strong an effect on the observed RRA as the bias observed due to chloroplast richness. Sometimes the two biases studied push in the same direction, as in *Populus tremula* (Ptr), which has a high chloroplast concentration and a high amplifiability, or *Capsella bursa-pastoris* (Cbe), which combines both a low chloroplast concentration and a low amplifiability (Fig. 4). Sometimes, by chance, both biases partially compensate, as in *Salvia pratensis* (Spr).

Even if the abundances observed by traditional surveys and those of metabarcoding reads are correlated (Yoccoz et al., 2012), it is necessary to be cautious when analyzing DNA metabarcoding data in terms of quantitative information. If we consider the estimation of biodiversity indices, the worst situation is the estimation of *α*-diversity. Because of all the biases acting simultaneously on DNA metabarcoding measures, but their good reproducibility, the information they provide is inherently relative. Relative in terms of abundances, DNA metabarcoding can at best provide relative abundances, but also relative because the values provided are biased. Therefore, only changes between measures are truly meaningful. Although it has been shown that *α*-diversity of plant communities can be correctly estimated from DNA metabarcoding data (Calderón-Sanou, Münkemüller, Boyer, Zinger, & Thuiller, 2020), the limited condition under which this is true, Hill numbers computed for *q* = 1, indicates that this is because at this level of weighting of rare versus abundant species by chance most of the biases are compensated. This phenomenon can also be observed in our results (Table 3), where ^1^*D* and ^2^*D* values estimated from raw RRA and corrected abundances do not strongly differ, while the error between RRA and theoretical composition decreases by a factor of two when using corrected abundances. This discrepancy between the decrease in error due to the correction and the not so good increase in the quality of the *α*-diversity estimates can be at least partially explained in **ℳ**_***G***_ by the abundances of the two most abundant species, *Briza media* (Bme) and *Rosa canina* (Rca), which have inverted abundances when estimated from RRA. For any study analyzing changes in diversity across time or ecological gradients, because metabarcoding measures are biased but accurate, the true *β*-diversity patterns can be easily detected using metabarcoding. In fact, because the biases are repeatable between measures, they often amplify the pattern because the errors correlate with the ecological signal. The problem of all these biases only arises when trying to disentangle the observed pattern from changes in specific species. Therefore, we can strongly encourage people to be very cautious when interpreting the observed pattern, and to be careful not to over-interpret changes in the abundance of a few species in the community as an ecological cause.

## Conclusion

We investigated two of the biases that prevent proper quantification of relative eDNA abundances in metabarcoding data. Despite their importance, these biases are far from being corrected or even considered in most current studies. In this study, we measure the two studied biases and propose a simple method to correct the amplification biases in the limit of extreme cases where some species are so strongly disadvantaged that they disappear from the raw results. The advantage of our method compared to the previous ones is that it is more robust to sample variability, while compared to the spiking-based method it does not require any change in metabarcoding protocols. This also allows the reanalysis of previously obtained results, providing the opportunity for a better ecological interpretation of them. By combining relative abundance correction and ddPCR to estimate the amount of target DNA in each sample, we can even consider the possibility of having access to an absolute quantification of DNA in the analyzed DNA extracts for each species instead of only relative abundances. This opens the possibility to increase the robustness of the quantitative interpretation of DNA metabarcoding results, although other biases still need to be assessed and modeled in a similar way to fully achieve the goal of truly quantitative metabarcoding.

## Acknowledgments

The authors thank Christian Miquel for the logistic support, Frédéric Boyer and Clément Lionnet for their help with the bioinformatics pipeline. This work was supported by the Alpalga project (ANR-20-CE02-0020).

## Data Accessibility and Benefit Sharing statement

### Data Accessibility statement

The data and analysis scripts are available on the project’s git page, https://github.com/LECA-MALBIO/metabar-bias.

### Authors’ contributions

SM, EC, CG and DP studied the PCR models. SM, EC, CG and PT designed the associated experimental protocol. SM, EC, CG and PT wrote the manuscript. DP contributed to the writing of the manuscript. EC and PT sampled the plants. DR and SM performed the extractions and metabarcoding PCRs. FL and SM performed the qPCR and ddPCR assays. SM wrote the analysis script. EC and CG supervised the project.

## References

Alberdi, A., & Gilbert, M.T.P. (2019, July). A guide to the application of Hill numbers to DNA-based diversity analyses. Molecular Ecology Resources, 19 (4), 804–817, 10.1111/1755-0998.13014

Andruszkiewicz Allan, E., Zhang, W.G., Lavery, A., Govindarajan, A. (2021, March). Environmental DNA shedding and decay rates from diverse animal forms and thermal regimes. Environmental DNA, 3 (2), 492–514, 10.1002/edn3.141

Barnosky, A.D., Matzke, N., Tomiya, S., Wogan, G.O.U., Swartz, B., Quental, T.B., … Ferrer, E.A. (2011, March). Has the Earth’s sixth mass extinction already arrived? Nature, 471 (7336), 51–57, 10.1038/nature09678 Retrieved from http://dx.doi.org/10.1038/nature09678

Beng, K.C., & Corlett, R.T. (2020, June). Applications of environmental DNA (eDNA) in ecology and conservation: opportunities, challenges and prospects. Biodiversity and Conservation, 29 (7), 2089–2121, 10.1007/s10531-020-01980-0

Bohmann, K., Evans, A., Gilbert, M.T.P., Carvalho, G.R., Creer, S., Knapp, M., … de Bruyn, M. (2014, June). Environmental DNA for wildlife biology and biodiversity monitoring. Trends in Ecology & Evolution, 29 (6), 358–367, 10.1016/j.tree.2014.04.003

Bou Dagher-Kharrat, M., Abdel-Samad, N., Douaihy, B., Bourge, M., Fridlender, A., Siljak-Yakovlev, S., Brown, S.C. (2013, December). Nuclear DNA C-values for biodiversity screening: Case of the Lebanese flora. Plant Biosystems - An International Journal Dealing with all Aspects of Plant Biology, 147 (4), 1228–1237, 10.1080/11263504.2013.861530 Retrieved from https://doi.org/10.1080/11263504.2013.861530

Boyer, F., Mercier, C., Bonin, A., Le Bras, Y., Taberlet, P., Coissac, E. (2016, January). obitools : a unix-inspired software package for DNA metabarcoding. Molecular Ecology Resources, 16 (1), 176–182, 10.1111/1755-0998.12428

Calderón-Sanou, I., Münkemüller, T., Boyer, F., Zinger, L., Thuiller, W. (2020, January). From environmental DNA sequences to ecological conclusions: How strong is the influence of methodological choices? Journal of biogeography, 47 (1), 193–206, 10.1111/jbi.13681 Retrieved from https://onlinelibrary.wiley.com/doi/abs/10.1111/jbi.13681

Carr, A.C., & Moore, S.D. (2012, May). Robust Quantification of Polymerase Chain Reactions Using Global Fitting. PLoS ONE, 7 (5), e37640, 10.1371/journal.pone.0037640 Retrieved 2023-07-12, from https://dx.plos.org/10.1371/journal.pone.0037640

Clarke, L.J., Soubrier, J., Weyrich, L.S., Cooper, A. (2014, November). Environmental metabarcodes for insects: in silico PCR reveals potential for taxonomic bias. Molecular Ecology Resources, 14 (6), 1160–1170, 10.1111/1755-0998.12265

Dopheide, A., Xie, D., Buckley, T.R., Drummond, A.J., Newcomb, R.D. (2019, January). Impacts of DNA extraction and PCR on DNA metabarcoding estimates of soil biodiversity. Methods in Ecology and Evolution, 10 (1), 120–133, 10.1111/2041-210X.13086 Retrieved 2023-07-19, from https://onlinelibrary.wiley.com/doi/10.1111/2041-210X.13086

Doyle, J.J. (1990). Isolation of plant dna from fresh tissue. Retrieved from https://api.semanticscholar.org/CorpusID:85677467

Elbrecht, V., & Leese, F. (2015, July). Can DNA-Based Ecosystem Assessments Quantify Species Abundance? Testing Primer Bias and Biomass—Sequence Relationships with an Innovative Metabarcoding Protocol. PLOS ONE, 10 (7), e0130324, 10.1371/journal.pone.0130324

Elbrecht, V., Peinert, B., Leese, F. (2017, September). Sorting things out: Assessing effects of unequal specimen biomass on DNA metabarcoding. Ecology and Evolution, 7 (17), 6918–6926, 10.1002/ece3.3192

Ficetola, G.F., & Taberlet, P. (2023, February). Towards exhaustive community ecology via dna metabarcoding. Molecular Ecology, mec.16881, 10.1111/mec.16881

Garrido-Sanz, L., Senar, M.A., Piñol, J. (2022, January). Relative species abundance estimation in artificial mixtures of insects using mitometagenomics and a correction factor for the mitochondrial DNA copy number. Molecular Ecology Resources, 22 (1), 153–167, 10.1111/1755-0998.13464

Gill, P., Bleka, O., Fonneløp, A.E. (2022, November). Limitations of qPCR to estimate DNA quantity: An RFU method to facilitate inter-laboratory comparisons for activity level, and general applicability. Forensic Science International: Genetics, 61, 102777, 10.1016/j.fsigen.2022.102777

Golczyk, H., Greiner, S., Wanner, G., Weihe, A., Bock, R., Börner, T., Herrmann, R.G. (2014, April). Chloroplast DNA in Mature and Senescing Leaves: A Reappraisal. The Plant Cell, 26 (3), 847–854, 10.1105/tpc.113.117465

Gold, Z., Shelton, A.O., Casendino, H.R., Duprey, J., Gallego, R., Van Cise, A., … Kelly, R.P. (2023, May). Signal and noise in metabarcoding data. PLOS ONE, 18 (5), e0285674, 10.1371/journal.pone.0285674 Retrieved 2023-07-23, from https://dx.plos.org/10.1371/journal.pone.0285674

Gottschalk, P.G., & Dunn, J.R. (2005, August). The five-parameter logistic: A characterization and comparison with the four-parameter logistic. Analytical Biochemistry, 343 (1), 54–65, 10.1016/j.ab.2005.04.035 Retrieved 2023-07-12, from https://linkinghub.elsevier.com/retrieve/pii/S0003269705003313

Hayward, A. (1998, June). Modeling and analysis of competitive RT-PCR. Nucleic Acids Research, 26 (11), 2511–2518, 10.1093/nar/26.11.2511

Hill, M.O. (1973, March). Diversity and Evenness: A Unifying Notation and Its Consequences. Ecology, 54 (2), 427–432, 10.2307/1934352

Kelly, R.P., Shelton, A.O., Gallego, R. (2019, December). Understanding PCR Processes to Draw Meaningful Conclusions from Environmental DNA Studies. Scientific Reports, 9 (1), 12133, 10.1038/s41598-019-48546-x

Kembel, S.W., Wu, M., Eisen, J.A., Green, J.L. (2012, October). Incorporating 16S Gene Copy Number Information Improves Estimates of Microbial Diversity and Abundance. PLoS Computational Biology, 8 (10), e1002743, 10.1371/journal.pcbi.1002743 Retrieved 2023-08-01, from https://dx.plos.org/10.1371/journal.pcbi.1002743

Klymus, K.E., Marshall, N.T., Stepien, C.A. (2017, May). Environmental DNA (eDNA) metabarcoding assays to detect invasive invertebrate species in the Great Lakes. PLOS ONE, 12 (5), e0177643, 10.1371/journal.pone.0177643

Krehenwinkel, H., Fong, M., Kennedy, S., Huang, E.G., Noriyuki, S., Cayetano, L., Gillespie, R. (2018, January). The effect of DNA degradation bias in passive sampling devices on metabarcoding studies of arthropod communities and their associated microbiota. PLOS ONE, 13 (1), e0189188, 10.1371/journal.pone.0189188

Krehenwinkel, H., Wolf, M., Lim, J.Y., Rominger, A.J., Simison, W.B., Gillespie, R.G. (2017, December). Estimating and mitigating amplification bias in qualitative and quantitative arthropod metabarcoding. Scientific Reports, 7 (1), 17668, 10.1038/s41598-017-17333-x

Lamb, P.D., Hunter, E., Pinnegar, J.K., Creer, S., Davies, R.G., Taylor, M.I. (2019, January). How quantitative is metabarcoding: A meta-analytical approach. Molecular Ecology, 28 (2), 420–430, 10.1111/mec.14920

Luo, M., Ji, Y., Warton, D., Yu, D.W. (2022, August). Extracting abundance information from dna-based data. Molecular Ecology Resources, 1755–0998.13703, 10.1111/1755-0998.13703

Matesanz, S., Pescador, D.S., Pías, B., Sánchez, A.M., Chacón-Labella, J., Illuminati, A., … Escudero, A. (2019, September). Estimating belowground plant abundance with DNA metabarcoding. Molecular Ecology Resources, 19 (5), 1265–1277, 10.1111/1755-0998.13049

Mehra, S., & Hu, W.-S. (2005, September). A kinetic model of quantitative real-time polymerase chain reaction. Biotechnology and Bioengineering, 91 (7), 848–860, 10.1002/bit.20555

Milivojević, T., Rahman, S.N., Raposo, D., Siccha, M., Kucera, M., Morard, R. (2021, October). High variability in SSU rDNA gene copy number among planktonic foraminifera revealed by single-cell qPCR. ISME Communications, 1 (1), 63, 10.1038/s43705-021-00067-3 Retrieved 2023-10-02, from https://www.nature.com/articles/s43705-021-00067-3

Moinard, S., Oudet, E., Piau, D., Coissac, E., Gonindard-Melodelima, C. (2022). The Fixed Landscape Inference MethOd (flimo): a versatile alternative to Approximate Bayesian Computation, faster by several orders of magnitude. arXiv,, 10.48550/ARXIV.2210.06520 (Publisher: arXiv Version Number: 3)

Mächler, E., Walser, J., Altermatt, F. (2021, July). Decision-making and best practices for taxonomy-free environmental DNA metabarcoding in biomonitoring using Hill numbers. Molecular Ecology, 30 (13), 3326–3339, 10.1111/mec.15725

Newton, C.R., & Graham, A. (2000). PCR (2. ed., repr ed.). Oxford: BIOS Scientific Publ.

Nichols, R.V., Vollmers, C., Newsom, L.A., Wang, Y., Heintzman, P.D., Leighton, M., … Shapiro, B. (2018, September). Minimizing polymerase biases in metabarcoding. Molecular Ecology Resources, 18 (5), 927–939, 10.1111/1755-0998.12895

Pawluczyk, M., Weiss, J., Links, M.G., Egaña Aranguren, M., Wilkinson, M.D., Egea-Cortines, M. (2015, March). Quantitative evaluation of bias in PCR amplification and next-generation sequencing derived from metabarcoding samples. Analytical and Bioanalytical Chemistry, 407 (7), 1841–1848, 10.1007/s00216-014-8435-y

Piñol, J., Mir, G., Gomez-Polo, P., Agustí, N. (2015, July). Universal and blocking primer mismatches limit the use of high-throughput DNA sequencing for the quantitative metabarcoding of arthropods. Molecular Ecology Resources, 15 (4), 819–830, 10.1111/1755-0998.12355

Pompanon, F., Deagle, B.E., Symondson, W.O.C., Brown, D.S., Jarman, S.N., Taberlet, P. (2012, April). Who is eating what: diet assessment using next generation sequencing: NGS DIET ANALYSIS. Molecular Ecology, 21 (8), 1931–1950, 10.1111/j.1365-294X.2011.05403.x

Pornon, A., Escaravage, N., Burrus, M., Holota, H., Khimoun, A., Mariette, J., … Andalo, C. (2016, June). Using metabarcoding to reveal and quantify plant-pollinator interactions. Scientific Reports, 6 (1), 27282, 10.1038/srep27282

Rao, X., Lai, D., Huang, X. (2013, September). A New Method for Quantitative Real-Time Polymerase Chain Reaction Data Analysis. Journal of Computational Biology, 20 (9), 703–711, 10.1089/cmb.2012.0279 Retrieved 2023-11-28, from http://www.liebertpub.com/doi/10.1089/cmb.2012.0279

Sakamoto, W., & Takami, T. (2018, June). Chloroplast DNA Dynamics: Copy Number, Quality Control and Degradation. Plant and Cell Physiology, 59 (6), 1120–1127, 10.1093/pcp/pcy084

Shelton, A.O., Gold, Z.J., Jensen, A.J., D’Agnese, E., Andruszkiewicz Allan, E., Van Cise, A., Kelly, R.P. (2022, November). Toward quantitative metabarcoding. Ecology,, 10.1002/ecy.3906

Sidstedt, M., Rådström, P., Hedman, J. (2020, April). PCR inhibition in qPCR, dPCR and MPS—mechanisms and solutions. Analytical and Bioanalytical Chemistry, 412 (9), 2009–2023, 10.1007/s00216-020-02490-2

Siljak-Yakovlev, S., Pustahija, F., Oli, E.M., Boguni, F., Muratovi, E., Bai, N., … Brown, S.C. (2010). Towards a Genome Size and Chromosome Number Database of Balkan Flora: C-Values in 343 Taxa with Novel Values for 242. Advanced science letters, 3 (2), 190–213, 10.1166/asl.2010.1115 Retrieved from https://www.ingentaconnect.com/content/asp/asl/2010/00000003/00000

Silverman, J.D., Bloom, R.J., Jiang, S., Durand, H.K., Dallow, E., Mukherjee, S., David, L.A. (2021, July). Measuring and mitigating PCR bias in microbiota datasets. PLOS Computational Biology, 17 (7), e1009113, 10.1371/journal.pcbi.1009113

Smets, W., Leff, J.W., Bradford, M.A., McCulley, R.L., Lebeer, S., Fierer, N. (2016). A method for simultaneous measurement of soil bacterial abundances and community composition via 16S rRNA gene sequencing. Soil Biology and Biochemistry, 96, 145–151, 10.7287/peerj.preprints.1318v1 Retrieved from https://peerj.com/preprints/1318

Svec, D., Tichopad, A., Novosadova, V., Pfaffl, M.W., Kubista, M. (2015, March). How good is a PCR efficiency estimate: Recommendations for precise and robust qPCR efficiency assessments. Biomolecular Detection and Quantification, 3, 9–16, 10.1016/j.bdq.2015.01.005

Taberlet, P., Bonin, A., Zinger, L., Coissac, E. (2018). Environmental DNA (Vol. 1). Oxford University Press.

Taberlet, P., Coissac, E., Pompanon, F., Brochmann, C., Willerslev, E. (2012, April). Towards next-generation biodiversity assessment using DNA metabarcoding: NEXT-GENERATION DNA METABARCODING. Molecular Ecology, 21 (8), 2045–2050, 10.1111/j.1365-294X.2012.05470.x

Taberlet, P., Coissac, E., Pompanon, F., Gielly, L., Miquel, C., Valentini, A., … Willerslev, E. (2007, January). Power and limitations of the chloroplast trnL (UAA) intron for plant DNA barcoding. Nucleic Acids Research, 35 (3), e14–e14, 10.1093/nar/gkl938

Thomas, A.C., Deagle, B.E., Eveson, J.P., Harsch, C.H., Trites, A.W. (2016, May). Quantitative DNA metabarcoding: improved estimates of species proportional biomass using correction factors derived from control material. Molecular Ecology Resources, 16 (3), 714–726, 10.1111/1755-0998.12490 Retrieved 2023-06-16, from https://onlinelibrary.wiley.com/doi/10.1111/1755-0998.12490

Ushio, M., Murakami, H., Masuda, R., Sado, T., Miya, M., Sakurai, S., … Kondoh, M. (2018). Quantitative monitoring of multispecies fish environmental dna using high-throughput sequencing. Metabarcoding and Metagenomics, 2, e23297, 10.3897/mbmg.2.23297 Retrieved from https://doi.org/10.3897/mbmg.2.23297 https://arxiv.org/abs/https://doi.org/10.3897/mbmg.2.23297

Valentini, A., Miquel, C., Nawaz, M.A., Bellemain, E., Coissac, E., Pompanon, F., … Taberlet, P. (2009). New perspectives in diet analysis based on DNA barcoding and parallel pyrosequencing: the trnL approach. Molecular Ecology Resources, 9, 51–60,

van der Loos, L.M., & Nijland, R. (2021, July). Biases in bulk: DNA metabarcoding of marine communities and the methodology involved. Molecular Ecology, 30 (13), 3270–3288, 10.1111/mec.15592 Retrieved 2022-08-31, from https://onlinelibrary.wiley.com/doi/10.1111/mec.15592

Wilder, M.L., Farrell, J.M., Green, H.C. (2023, March). Estimating edna shedding and decay rates for muskellunge in early stages of development. Environmental DNA, 5 (2), 251–263, 10.1002/edn3.349

Willerslev, E., Davison, J., Moora, M., Zobel, M., Coissac, E., Edwards, M.E., … Taberlet, P. (2014, February). Fifty thousand years of Arctic vegetation and megafaunal diet. Nature, 506 (7486), 47–51, 10.1038/nature12921

Yang, C., Bohmann, K., Wang, X., Cai, W., Wales, N., Ding, Z., Yu, D.W. (2021, July). Biodiversity Soup II: A bulk-sample metabarcoding pipeline emphasizing error reduction. Methods in Ecology and Evolution, 12 (7), 1252–1264, 10.1111/2041-210X.13602

Yoccoz, N.G., Bråthen, K.A., Gielly, L., Haile, J., Edwards, M.E., Goslar, T., … Taberlet, P. (2012, August). DNA from soil mirrors plant taxonomic and growth form diversity: DNA FROM SOIL MIRRORS PLANT DIVERSITY. Molecular Ecology, 21 (15), 3647–3655, 10.1111/j.1365-294X.2012.05545.x

Zoschke, R., Liere, K., Börner, T. (2007, April). From seedling to mature plant: Arabidopsis plastidial genome copy number, RNA accumulation and transcription are differentially regulated during leaf development: Plastome copy number in Arabidopsis leaf development. The Plant Journal, 50 (4), 710–722, 10.1111/j.1365-313X.2007.03084.x

