## Supporting information for "Towards quantitative DNA Metabarcoding: A method to overcome PCR amplification bias"

### Supporting information for online publication

| Species | Barcode | 1C value (pg) |
|---|---|---|
| *Briza media* (Bme) | atccgtgttttgagaaaacaagggggttctcgaa<br>ctagaatacaaaggaaaag | 6.35[1] |
| *Rosa canina* (Rca) | atcccgttttatgaaaacaaacaaggtttcagaa<br>agcgagaataaataaag | 1.40 |
| *Lotus corniculatus* (Lco) | atcctgctttacgaaaacaagggaaagttcagtt<br>aagaaagcgacgagaaaaatg | 0.87[1] |
| *Populus tremula* (Ptr) | atcctatttttcgaaaacaaacaaaaaaacaaac<br>aaaggttcataaagacagaataagaatacaaaag | 0.45 |
| *Salvia pratensis* (Spr) | atcctgttttctcaaaacaaaggttcaaaaaacg<br>aaaaaaaaaaag | 0.46 |
| *Lonicera xylosteum* (Lxy) | atccagttttccgaaaacaagggtttagaaagca<br>aaaatcaaaaag | 0.70 |
| ***Fraxinus excelsior* (Fex)** | **atcctgttttcccaaaacaaaggttcagaaagaa<br>aaaag** | **0.84** |
| *Acer campestre* (Aca) | atcctgttttacgagaataaaacaaagcaaacaa<br>gggttcagaaagcgagaaaggg | 1.02[1] |
| ***Capsella bursa-pastoris* (Cbp)** | **atcctggtttacgcgaacacaccggagtttacaa<br>agcgagaaaaaagg** | **0.40** |
| *Geranium robertianum* (Gro) | atcctttttttacgaaaataaagaggggctcacaa<br>agcgagaatagaaaaaaag | 1.76[2] |
| ***Carpinus betulus* (Cbe)** | **atcctgttttcccaaaacaaataaaacaaattta<br>agggttcataaagcgagaataaaaaag** | **1.03** |
| *Abies alba* (Aal) | atccggttcatagagaaaagggtttctctccttc<br>tcctaaggaaagg | 17.29 |
| *Rhododendron ferrugineum* (Rfe) | atccttttttcgcaaacaaacaaagattccgaaa<br>gctaaaaaaaag | 0.74[2] |

**Supplementary Table 1**: Metabarcodes and genome sizes of the plants used for the three mock communities for the *Sper01* marker. In our data, the sequence of *Salvia pratensis* has an insertion (a, position 35) compared to the reference sequence. 1C value caracterises the genome size and can be found in the Kew database (https://cvalues.science.kew.org/). [1]: average value of different assays. [2]: average C-value of species in same genus when species C-value is missing.

| Species | Concentration ($ng/\mu l$) | | | Copies per well ($2\mu l$ of DNA) | | | Rank |
|---|---|---|---|---|---|---|---|
| | $\mathcal{M}_U$ | $\mathcal{M}_T$ | $\mathcal{M}_G$ | $\mathcal{M}_U$ | $\mathcal{M}_T$ | $\mathcal{M}_G$ | $\mathcal{M}_G$ |
| *Briza media* | 0.076 | 0.038 | $4.9 \times 10^{-1}$ | $1.9 \times 10^4$ | $9.7 \times 10^3$ | $1.2 \times 10^5$ | 1 |
| *Rosa canina* | 0.045 | 0.038 | $1.5 \times 10^{-1}$ | $1.9 \times 10^4$ | $1.6 \times 10^4$ | $6.2 \times 10^4$ | 2 |
| *Lotus corniculatus* | 0.068 | 0.038 | $1.1 \times 10^{-1}$ | $1.9 \times 10^4$ | $1.1 \times 10^4$ | $3.1 \times 10^4$ | 3 |
| *Populus tremula* | 0.038 | 0.038 | $3.1 \times 10^{-2}$ | $1.9 \times 10^4$ | $1.9 \times 10^4$ | $1.6 \times 10^4$ | 4 |
| *Salvia pratensis* | 0.062 | 0.038 | $2.5 \times 10^{-2}$ | $1.9 \times 10^4$ | $1.2 \times 10^4$ | $7.8 \times 10^3$ | 5 |
| *Lonicera xylosteum* | 0.13 | 0.038 | $2.6 \times 10^{-2}$ | $1.9 \times 10^4$ | $5.7 \times 10^3$ | $3.9 \times 10^3$ | 6 |
| ***Fraxinus excelsior*** | **0.11** | **0.038** | $\mathbf{1.1 \times 10^{-2}}$ | $\mathbf{1.9 \times 10^4}$ | $\mathbf{6.7 \times 10^3}$ | $\mathbf{1.9 \times 10^3}$ | **7** |
| *Acer campestre* | 0.071 | 0.038 | $3.6 \times 10^{-3}$ | $1.9 \times 10^4$ | $1.0 \times 10^4$ | $9.7 \times 10^2$ | 8 |
| ***Capsella bursa-pastoris*** | **0.25** | **0.038** | $\mathbf{6.3 \times 10^{-3}}$ | $\mathbf{1.9 \times 10^4}$ | $\mathbf{3.0 \times 10^3}$ | $\mathbf{4.8 \times 10^2}$ | **9** |
| *Geranium robertianum* | 0.12 | 0.038 | $1.2 \times 10^{-3}$ | $1.9 \times 10^4$ | $6.3 \times 10^3$ | $2.4 \times 10^2$ | 10 |
| ***Carpinus betulus*** | **0.18** | **0.038** | $\mathbf{1.2 \times 10^{-3}}$ | $\mathbf{1.9 \times 10^4}$ | $\mathbf{4.1 \times 10^3}$ | $\mathbf{1.2 \times 10^2}$ | **11** |
| *Abies alba* | 0.11 | 0.038 | $3.4 \times 10^{-4}$ | $1.9 \times 10^4$ | $6.9 \times 10^3$ | $6.1 \times 10^1$ | 12 |
| *Rhododendron ferrugineum* | 0.25 | 0.038 | $4.0 \times 10^{-4}$ | $1.9 \times 10^4$ | $2.9 \times 10^3$ | $3.0 \times 10^1$ | 13 |

**Supplementary Table 2**: Total DNA concentrations (in the samples) and number of molecules per well of plants used for the three mock communities. The rank for $\mathcal{M}_G$ stands for the decreasing abundance in terms of target DNA.

| Species | Probe name | Probe sequence | Positions | Tm (salt-adjusted) |
|---|---|---|---|---|
| *Fraxinus excelsior* | Fex | ttttcccaaaacaaaggttcagaaagaaaa | 7 - 36 | 63.9°C |
| *Capsella bursa-pastoris* | Cbp | aacacaccggagtttacaaagcgag | 16 - 40 | 65.8°C |
| *Carpinus betulus* | CbeA | tcctgttttcccaaaacaaataaaacaaat | 1 - 30 | 62.5°C |
| | CbeB | ttaagggttcataaagcgagaataaaaaag | 32 - 61 | 63.9°C |

**Supplementary Table 3**: Taqman probes designed for three different species for the *Sper01* marker. For *Carpinus betulus*, two probes were designed. The positions desginate the bases of the original metabarcodes.
